# EvolQG - An R package for evolutionary quantitative genetics

**DOI:** 10.1101/026518

**Authors:** Diogo Melo, Guilherme Garcia, Alex Hubbe, Ana Paula Assis, Gabriel Marroig

## Abstract

We present an open source package for performing evolutionary quantitative genetics analyses in the R environment for statistical computing. Evolutionary theory shows that evolution depends critically on the available variation in a given population. When dealing with many quantitative traits this variation is expressed in the form of a covariance matrix, particularly the additive genetic covariance matrix or sometimes the phenotypic matrix, when the genetic matrix is unavailable. Given this mathematical representation of available variation, the **EvolQG** package provides functions for calculation of relevant evolutionary statistics, estimation of sampling error, corrections for this error, matrix comparison via correlations and distances, and functions for testing evolutionary hypotheses on taxa diversification.

## Introduction

Quantitative genetics deals with the evolution and inheritance of continuous traits, like body size, bone lengths, gene expressions or any other inheritable characteristic that can be measured on a continuous scale, or which can be transformed to a continuous scale. This framework has been used in selective breeding and in describing the different sources of variation in natural populations, as well as understanding the interaction of evolutionary processes with this variation (Lynch and Walsh. 1998). Quantitative genetics has been successful in describing short term evolution, and is also useful in understanding diversification at a macroevolutionary level. The core development of modern evolutionary quantitative genetics started with the generalization of the univariate breeders equation to the multivariate response to selection equation, derived by Lande and also referred to as the Lande equation (Lande 1979; Roff 2003).

The Lande equation relates the evolutionary change in trait means of a given population (Δ*z*) to the interaction between the additive genetic variation (**G**-matrix) of this population and the directional selection (*β*) acting on this population. The additive genetic variation of a population is represented by a symmetric square matrix called the **G**-matrix, which contains the additive genetic variance of each trait on the diagonal and the additive genetic covariance between traits on the off-diagonal elements. From the Lande equation, Δ*z* = **G***β*, we can see that different populations may present markedly different responses (Δ*z*) to the same directional selection (*β*) simply because these populations have distinct **G**-matrices. Other evolutionary forces affecting populations are also influenced by available variation, e.g., based on the **G**-matrix it is possible to test if morphological differentiation of extant taxa is compatible with genetic drift or stabilizing selection (e.g., Ackermann and Cheverud (2004); Marroig and Cheverud (2004)). Thus, describing and understanding changes in standing variation among populations ((Cheverud 1982; Lofsvold 1986; Marroig and Cheverud 2001)) as well as understanding constraints imposed by populations standing variation (e.g., Schluter (1996); Marroig and Cheverud (2010); Hansen and Houle (2008); Porto *et al.* (2013)) are major elements in evolutionary quantitative genetics.

In this article we describe the **EvolQG** package, developed to deal with the evolutionary quantitative genetics questions addressed above in the R environment for statistical computing (R Core Team 2014). Our goal was to provide a suite of tools in a single consistent source, and to standardize and facilitate the adoption of these tools.

## Measurement error estimation

Before estimating a **G**-matrix, it is important to evaluate the influence of measurement error in data collection, since measurement error can drastically bias further analyses (Falconer and Mackay 1996). Measurement error can be estimated by measuring each individual at least twice and estimating the amount of variation associated with each individual, which is the measurement error, in relation to total variation (i.e., the sum of within and between individuals variation) using an analysis of variance. The proportion of variance not associated with the individuals is called the repeatability (Lessells and Boag 1987). A repeatability of 1 means that no variation is associated with measurement error. The function **CalcRepeatability()** performs the calculation described in Lessells and Boag (1987) for a set of multivariate traits measured at least twice for each individual.

## Matrix estimation

In the rest of this article we assume that the covariance matrix of interest has already been estimated by some procedure. This can be a simple covariance of all the observed traits, or an estimated parameter from a more complicated linear model. The simplest case of a linear model approach would be using a multivariate analysis of covariance (MANCOVA) to control for differences in trait means that are not of immediate interest in the analyses (e.g., sexual dimorphism, geographic variation, etc.). The residual pooled within-group covariance matrix can be used in subsequent analysis (Marroig and Cheverud 2001). The **EvolQG** package function **CalculateMatrix()** uses R’s **lm()** model object to calculate variance-covariance matrices adjusting for the proper degrees of freedom in a simple fixed-effects MANCOVA. Of course more complicated methods may be used to obtain **G**-matrices, such as an animal model or a mixed model (Lynch and Walsh. 1998; Runcie and Mukherjee 2013).

Accurate **G**-matrix estimation can be hard to achieve, requiring large sample sizes, many families and known genealogies (Steppan *et al.* 2002). One alternative that is usually more feasible is to use the phenotypic covariance matrix (the **P**-matrix) as a proxy of the population’s **G**-matrix (Cheverud 1988; Roff 1995). Conditions on where this approximation is reasonable depend on the structure of developmental and environmental effects, and testing for similarity is an empirical question that should be undertaken before using the **P**-matrix as a proxy for the **G**-matrix, ideally by direct comparison (e.g.,Garcia *et al.* (2014)). As a general rule, high similarity between populations’ **P**-matrices is a good indicator of high similarity between **P** and **G**, and of a stable shared **G**-matrix pattern, since the similarity between populations must come from either a common genetic structure, or the unlikely scenario of a different genetic structure buffered by an exactly compensating environmental structure in each population that leads to high similarity between phenotypic covariation.

Some of the methods described below are not applicable to covariance matrices, only to correlation matrices. Correlations are standardized measures of association that are bounded between [– 1, 1], and, unlike covariances, can be directly compared for pairs of traits with different scales. In most instances, correlation matrices can be obtained directly from covariance matrices by using the R function **cov2cor()**.

## Matrix error and repeatabilities

A **G**-matrix will always be estimated with error (Hill and Thompson 1978; Meyer and Kirkpatrick 2008; Marroig *et al.* 2012), and it is important to take this error into account in further analyses. In some circumstances we want to compare two or more **G**-matrices, calculating the matrices correlations (see section **Matrix Comparison**). However, due to error in estimating these matrices, their correlations will never be one, even if the actual population parameter values are identical (Cheverud 1988). Thus, matrix repeatabilities are used to correct matrix correlations by taking sampling error into account. The basic premise of all the methods is that taking repeated samples from the same population and comparing the resulting matrices would still give correlations that are lower than 1. We estimate the maximum correlation between matrices taken from the same population and correct the observed correlation by this maximum value. The corrected correlation between two observed matrices will be given by the original correlation divided by the geometric mean of their repeatabilities. If the repeatability of both matrices is one, the observed correlation does not change under the correction, and lower repeatabilities yield larger corrections. A number of methods for repeatability estimation are provided, and their results can be passed on to the functions that calculate matrix correlations (section **Matrix Comparison**):

**AlphaRep():** Cheverud (1988) describes an analytical expression for the repeatability of a correlation matrix. This expression is asymptotically convergent, so it should be used only when sample sizes are large, at least larger than the number of traits.

**BootstrapRep():** We may estimate the repeatability of the covariance (or correlation) structure of a given data set using a bootstrap procedure, sampling individuals with replacement from the data set and calculating a covariance (or correlation) matrix from each sample. The mean value of the correlation between the random sample matrix and the original estimated matrix is an estimate of the repeatability. This method has the advantage of not assuming any distribution on the data, but does provide inflated estimates of repeatabilities for small data sets. Even so, upwardly biased matrix repeatabilities are not so problematic, since they lead to conservative corrections of matrix correlations. However, users should be aware of this bias and should not interpret a high repeatability obtained from a small data set as indicating that the parameter is well estimated.

**MonteCarloRep():** We can use the observed covariance (or correlation) matrix as the Σ parameter in a multivariate normal distribution, and produce samples from this distribution, using a fixed sample size. The covariance (or correlation) matrix for each sample is compared to the observed matrix, and the mean of these comparisons is an estimate of the repeatability (Marroig and Cheverud 2001). This method has the advantage of being easy to apply to matrices coming from linear models with many controlled effects, and not requiring the original data.

Sometimes the question we are trying to answer does not involve matrix comparisons, so other methods of assessing and correcting for error are needed.

**Rarefaction():** Rarefaction consists of taking progressively smaller samples with replacement from the original data set, calculating some statistic on each data set and comparing this with the full data set. This gives a general idea of how the inferences would change if we had smaller sample sizes, and how robust our data set is with respect to sampling. The default operation is to calculate the covariance or correlation matrices and compare them using any of the matrix comparison methods (see section **Matrix Comparison**).

**ExtendMatrix():** Marroig *et al.* (2012) showed that sampling error on covariance matrix estimation can have a dramatic effect on the reconstruction of net selection gradients using the multivariate response to selection equation (Lande 1979). One way to improve estimates is the simple procedure of “extending” the eigenvalues of the covariance matrix, where all the eigenvalues lower than a certain threshold are substituted by the smallest eigenvalue above the threshold. This causes minimal changes in the distribution of phenotypes, but improves dramatically the estimates of net selection gradients. See Marroig et al. (2012) for a thorough examination of the performance and consequences of the extension method on simulated and real data sets.

## Evolutionary statistics

Hansen and Houle (2008) provide a suite of statistics that have fairly good biological interpretations for a given **G**- or **P**-matrix. Marroig *et al.* (2009) is a comprehensive example of how these statistics may be used for interpreting morphological data.

The function **MeanMatrixStatistics()** calculates most of these statistics and their distributions, as shown below.

In the following, *E*[·]*β* represents the expected value over many random *β* vectors with unit norm, *<*·, ·*>*represents the dot product between two vectors, *cos*(·,·) is the cosine between two vectors, **G** is an arbitrary covariance matrix, **G**-1 is the inverse **G**, *tr*(**G**) is the trace of **G**, and **‖ · ‖** the Euclidean norm of a vector. **MeanMatrixStatistics()** calculates:

- Mean aquared correlation (*r*^2^): Given a correlation matrix, the elements below the diagonal are squared and averaged resulting in a measure of integration, that is, overall association between traits (also see the section **Modularity and Integration** and Pavlicev *et al.* (2009)).
- Coefficient of variation of eigenvalues (ICV): A measure of integration that is suitable for covariance matrices, as it takes the amount of variation into account. Notice that at least for mammals, mean squared correlations and ICV generally have very high correlation, but can lead to different conclusions if the traits included in the analysis have very different variances (due to scale, for example). If *σλ* is the standard deviation of the eigenvalues of a covariance matrix, and 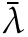 is the mean of the eigenvalues, the ICV is:

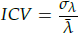
- Percent of variation in first principal component: If 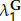 is the leading eigenvalue of **G**, we calculate this percentage as:

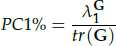
- Evolvability (Fig. 1): The mean projection of the response to random selection gradients with unit norm onto the selection gradient. This is a measure of a population’s available variation in the direction of a particular selection gradient, averaged across all directions (Hansen and Houle 2008).

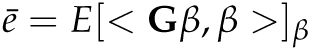
- Flexibility (Fig. 1): The mean cosine of the angle between random selection gradients and the corresponding responses. Flexibility measures on average how the response to selection aligns with the selection gradient (Marroig *et al.* 2009).

**Figure 1.**
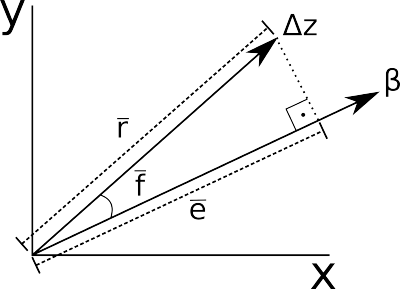
Graphical representation of evolvability (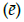), respondability (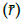) and flexibility (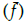) for a single selection gradient (*β*) and the corresponding response (Δ*z*) in the two dimensions defined by traits *x* and *y*.

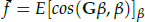
- Respondability (Fig. 1): Mean norm of the response to random selection gradients with unit norm. It also estimates how fast the population mean will change under directional selection (Hansen and Houle 2008; Marroig *et al.* 2009).

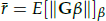
- Conditional Evolvability: Measures the mean response to selection in the direction of a given *β* when other directions are under stabilizing selection (Hansen and Houle 2008).

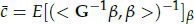
- Autonomy: Measures the proportion of variance in the direction of a given *β* that is independent from variation in other directions. Therefore, mean Autonomy can also be calculated as the mean ratio between Conditional Evolvability 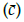 and Evolvability 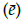 (Hansen and Houle 2008).

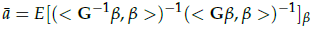
- Constraints: The mean correlation between the response vector to random selection gradients and the matrix’s first principal component (Marroig *et al.* 2009). If 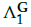 is the first principal component of **G**, constraints are measured as:

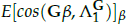

## Matrix comparison

A **G**-matrix describes how the variation in particular populations is structured, but frequently the relevant question is how similar or dissimilar two populations are with respect to this standing variation. Since no two populations are identical, different patterns of variation are the norm. Depending on the evolutionary question at hand, different methods of comparing variation may be required. One possible application of matrix comparisons is when we wish to apply the Lande equation to micro and macroevolution, since this requires some additional assumptions, such as a relative stability of the **G**-matrix over generations. Comparing extant covariance matrices is a test of this required stability (e.g. Marroig and Cheverud (2001)). For a thoughtful discussion on the biological relevance of statistical significance in matrix comparisons, see the discussion in Haber (2015).

### Matrix correlations

One approach to estimate the similarity or dissimilarity between matrices is to calculate the correlation between these matrices. **EvolQG** provides several functions for pairwise matrix correlation.

#### RandomSkewers()

The Random Skewers (RS) method makes use of the Lande equation (Lande 1979), Δ*z* = *Gβ*, where Δ*z* represents the vector of response to selection, **G** the **G**-matrix and *β* the directional selection vector, or selection gradient. In the RS method, the two matrices being compared are multiplied by a large number of normalized random selection vectors, and the resulting response vectors to the same selection vector are compared via a vector correlation (the cosine between the two vectors). The mean value of the correlation between the responses to the same selective pressure is used as a simple statistic of how often two populations respond similarly (in the same direction) to the same selective pressure:

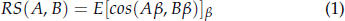

Where *E*[·]*β* is the expected value over random selection vectors *β*. Significance in the random skewers comparison can be determined using a null expectation of correlation between random vectors. If the observed correlation between two matrices is above the 95% percentile of the distribution of correlations between random vectors, we consider the correlation significant and infer that there is evidence the two populations behave similarly under directional selection. Other implementations of the RS method sometimes resort to other forms of calculating significance, such as generating random matrices and creating a random distribution of correlations between matrices. This is difficult to do because generating random matrices with the properties of biological covariance structures is hard, see the **RandomMatrix()** function for a quick discussion on this. The RS values range between -1 (the matrices have the opposite structure) and 1 (the matrices share the same structure), and zero means the matrices have distinct structures.

#### MantelCor()

Correlation matrices can be compared using a simple Pearson correlation between the corresponding elements. Significance of this comparison must take the structure into account, so it is calculated by a permutation scheme, in which a null distribution is generated by permutation of rows and columns in one of the matrices and repeating the element-by-element correlation. The observed correlation is significant when it is larger than the 95% quantile of the permuted distribution. This method can not be used on covariance matrices because the variances might be very different, leading to large differences in the scale of the covariances. This scale difference can lead to a massive inflation in the correlation between matrices. The correlation between matrices range between -1 (the matrices have the opposite structure) and 1 (the matrices share the same structure), and zero means the matrices have distinct structures.

#### KzrCor()

The Krzanowski shared space, or Krzanowski correlation, measures the degree to which the first principal components (eigenvectors) span the same subspace (Krzanowski 1979; Aguirre *et al.* 2014), and is suitable for covariance or correlation matrices. If two *n × n* matrices are being compared, the first 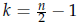 principal components from one matrix are compared to the first *k* principal components of the other matrix using the square of the vector correlations, and the sum of the correlations is a measure of how congruent the spanned subspaces are. We can write the Krzanowski correlation in terms of the matrices’ principal components (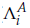 being the i*th* principal component of matrix *A*):

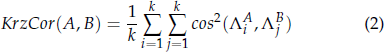

The Krzanowski correlation values range between 0 (two subspaces are dissimilar) and 1 (two subspaces are identical).

#### PCAsimilarity()

The Krzanowski correlation compares only the subspace shared by roughly the first half of the principal components, but does not consider the amount of variation each population has in these directions of the morphological space (Yang and Shahabi 2004). In order to take the variation into account, we can add the eigenvalue associated with each principal component into the calculation, effectively pondering each correlation by the variance in the associated directions. If 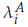 is the i*th* eigenvalue of matrix A, we have:

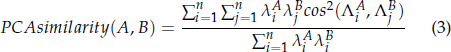

Note the sum spans all the principal components, not just the first *k* as in the Krzanowski correlation method. This method gives correlations that are very similar to the RS method, but is much faster. The PCA similarity values range between 0 (the shared subspaces have no common variation) and 1 (the shared subspaces have identical variation).

#### SRD()

The RS method can be extended to give information into which traits contribute to differences in terms of the pattern of correlated selection due to covariation between traits in two populations (Marroig *et al.* 2011). The Selection Response Decomposition does this by treating the terms of correlated response in the Lande equation as separate entities. Writing out the terms in the multivariate response to selection equation:

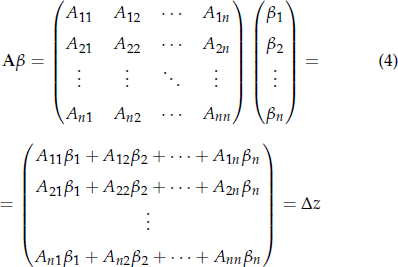

Separating the terms in the sums of the right hand side:

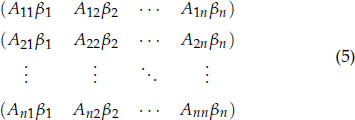

Each of these row vectors 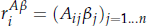 are the components of the response to the selection gradient *β* on trait *i*. The term *A_ii_ β_i_* represents the response to direct selection on trait *i*, and the terms (*A_ij_ β_j_*)*i/*=*j* represent the response to indirect selection due to correlation with the other traits. Given two matrices, *A* and *B*, we can measure how similar they are in their pattern of correlated selection on each trait by calculating the correlation between the vectors *r_i_* for each trait for random selection vectors of unit norm. The mean SRD score for trait *i* is then:

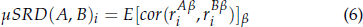

And the standard deviation of the correlations gives the variation in SRD scores:

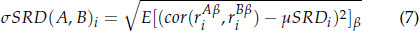

When the same trait in different matrices share a correlated response pattern, *µSRD* is high and *σSRD* is low; if the correlated response pattern is different, *µSRD* is low and *σSRD* is high. See (Marroig *et al.* 2011) for details and examples.

### Matrix distances

Another approach to estimate the similarity or dissimilarity between matrices is to calculate the distance between a pair of matrices. Matrices distances are different from correlations in that correlations are limited to [−1, +1], while distances must only be positive. Also, smaller values of distances mean more similarity. Two distances are in use in the current evolutionary literature, and are implemented in the function **MatrixDistance()**.

- Overlap distance: Ovaskainen *et al.* (2008) proposed a distance based on probability distributions, where two covariance matrices would have a distance proportional to how distinguishable they are. This distance is natural if we think of covariance matrices as describing the probability distribution of phenotypes or additive values in the population. The higher the probability of a random draw coming from the distribution defined by one of the matrices being misclassified as coming from the distribution defined by the other, the lower the distance. For two probability distributions *f* and *g*, the probability of mis-classifying a draw from *f* as coming from *g* is:

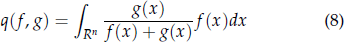

where *n* is the dimensionality of the space in which the distributions are defined. If the distributions are indistinguishable, *q*(*f*, *g*) = 1/2, if they are completely distinguishable *q*(*f*, *g*) = 0. We can then define the distance as:

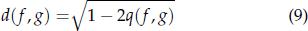 Since *q*(*f*, *g*) is symmetrical, *d*(*f*, *g*) is also symmetrical, and the square root guaranties that *d*(*f*, *g*) satisfies the triangle inequality (Ovaskainen *et al.* 2008). Calculation is straight forward and can be done with a simple sampling Monte Carlo scheme, see Ovaskainen *et al.* (2008) for details.
- Riemann distance: Mitteroecker and Bookstein (2009) use a Riemannian metric in the space of positive definite matrices (either covariance or correlation matrices), based on exponential mapping (Moakher 2006) to quantify transition in the ontogenetic trajectory of phenotypic covariance matrices. This metric is based on the eigenvalues of the product of one matrix to the inverse of the other. If *λ_i_* are the eigenvalues of *A*^−1^ *B* (or *AB*^−1^), we have:

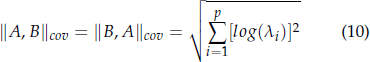 This distance has the advantage of being invariable under changes in the base used to represent the matrices. See Mitteroecker and Bookstein (2009) for a discussion on the biological relevance of this distance.

## Phylogenetic comparisons

**AncestralStates():** Ancestral state reconstruction of continuous traits using maximum likelihood can be performed in R (for example, using ape or phytools), but most packages don’t provide a simple interface for multivariate data. **AncestralStates()** is a wrapper for **fastAnc()** in phytools (Revell 2012) that works on multivariate data, reconstructing each character independently.

**PhyloW():** Given a set of covariance matrices for the terminal taxa in a phylogeny, we can estimate the covariance matrix for internal nodes by taking means over sister taxa, weighted by sample size. The mean matrix at the root node is the within-group covariance matrix in a MANCOVA with the terminal clades as the fixed effects. **PhyloW()** does this by taking a tree and a set of measurements (covariance matrices) and returns means for internal nodes.The implementation is generic, so this function can also be used to calculate weighted means for any numerical measurement with an addition operation implemented in R.

While using the within-group covariance matrix is a reasonable alternative as the estimator of an ancestral covariance matrix, this ignores branch lengths, and so should be used carefully when matrix differences are correlated to phylogenetic distance. An alternative when matrix evolution depends of branch lengths is to reconstruct every position of the covariance matrix independently via maximum likelihood, but this method can result in non positive-definite estimates.

**PhyloCompare():** Sometimes it is not practical to pair-wise compare every single population in a study, since for a large number of populations these results can be difficult to interpret. In these cases, comparing populations in a phylogeneticaly structured way can be helpful in detecting major transitions or differences between clades. **PhyloCompare()** takes estimates for all the nodes in a tree and compares sister groups by any comparison method, providing comparison values for every inner node.

## Hypothesis testing

### Modularity and integration

Modularity is a general concept in biology, and refers to a pattern of organization that is widespread in many biological systems. In modular systems, we find that some components of a given structure are more related or interact more between themselves than with other components. These highly related groups are termed modules. The nature of this interaction will depend on the components being considered, but may be of any kind, like physical contact between proteins, joint participation of enzymes in given biochemical pathways, or high correlation between quantitative traits in a population. This last kind of modularity is called variational modularity, and is characterized by high correlations between traits belonging to the same module and low correlation between traits in different modules (Wagner *et al.* 2007). In the context of morphological traits, variational modularity is associated with the concept of integration (Olson and Miller 1958), that is, the tendency of morphological systems to exhibit correlations due to common developmental factors and functional demands (Cheverud 1996; Hallgrímsson *et al.* 2009).

Both modularity and integration may have important evolutionary consequences, since sets of integrated traits will tend to respond to directional selection in an orchestrated fashion due to genetic correlations between them; if these sets are organized in a modular fashion, they will also respond to selection independently of one another (Marroig *et al.* 2009). At the same time, selection can alter existing patterns of integration and modularity, leading to traits becoming more or less correlated (Jones *et al.* 2004; Melo and Marroig 2015). The pattern of correlation between traits in a **G**-matrix then carries important information on the expected response to selection and on the history of evolutionary change of a given population.

**TestModularity():** Variational modularity can be assessed by comparing modularity hypothesis (derived from development and functional considerations) with the observed correlation matrix. If two traits are in the same variational module, we expect the correlation between them to be higher than between traits belonging to different modules. We test this by creating a modularity hypothesis matrix and comparing it via Mantel correlation with the observed correlation matrix. The modularity hypothesis matrix consists of binary matrix where each row and column corresponds to a trait. If the trait in row *i* is in the same module of the trait in column *j*, position (*i*, *j*) in the modularity hypothesis matrix is set to one, if these traits are not in the same module, position (*i*, *j*) is set to zero. Significant correlation between the hypothetical matrix representing a modularity hypothesis and the observed correlation matrix represents evidence of the existence of this variational module in the population. We also measure the ratio between correlations within a module (AVG+) and outside the module (AVG−). This ratio (AVG+/AVG−) is called the AVG Ratio, and measures the strength of the within-module association compared to the overall association for traits outside the module. The higher the AVG Ratio, the bigger the correlations within a module in relation to all other traits associations in the matrix (e.g., Porto *et al.* (2009)). **TestModularity()** also provides the Modularity Hypothesis Index, which is the difference between AVG+ and AVG-divided by the coefficient of variation of eigenvalues. Although the AVG Ratio is easier to interpret (how many times greater the within-module correlation is compared to the between-module correlation) than the Modularity Hypothesis Index, the AVG Ratio cannot be used when the observed correlation matrix presents correlations that differ in sign, and this is usually the case for residual matrices after size removal (for example with **RemoveSize()**, but see Jungers *et al.* (1995) for other alternatives). In these cases, Modularity Hypothesis Index is useful and allows comparing results between raw and residual matrices (Porto *et al.* 2013).

**LModularity():** If no empirical or theoretical information is available for creating modularity hypothesis, such as functional or developmental data, we can try to infer the modular partition of a given population by looking only at the correlation matrix and searching for the trait partition that minimizes some indicator of modularity. Borrowing from network theory, we can treat a correlation matrix as a fully connected weighted graph, and define a Newman-like modularity index (Newman 2006). If *A* is a correlation matrix we define *L* modularity as:

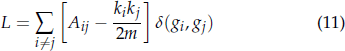

The terms *g_i_* and *g_j_* represent the partition of traits, that is, in what modules the traits *i* and *j* belong to. The function *δ*(·,·) is the Kronecker delta, where:

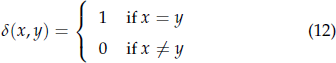

This means only traits in the same module contribute to the value of *L*. The term *k*_*i*_ represent the total amount of correlation attributed to trait *i*, or the sum of the correlation with trait *i*:

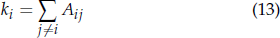

And *m* is the sum of all *k* (*m* = Σ_*i*_ *k*_*i*_). The term 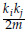 plays the role of a null expectation for the correlation between the traits *i* and *j*. This choice for the null expectation is natural when we impose that it must depend on the values of *k*_*i*_ and *k*_*j*_ and must be symmetrical (Newman 2006). So, traits in the same module with correlations higher than the null expectation will contribute to increase the value of *L*, while traits in the same module with correlation less than the null expectation will contribute to decrease *L*. With this definition of *L*, we use a Markov Chain Monte Carlo annealing method to find the partition of traits (values of *g*_*i*_) that maximizes *L*. This partition corresponds to the modularity hypothesis inferred from the correlation matrix, and the value of *L* is a measure of modularity comparable to the AVG Ratio. The igraph package (Csardi and Nepusz 2006) provides a number of similar community detection algorithms that can also be used in correlation matrices.

**RemoveSize():** If the first principal component of a covariance or correlation matrix corresponds to a very large portion of its variation, and all (or most) of the entries of the first principal component are of the same sign (a *size* principal component, see Marroig and Cheverud (2010)), it is useful to look at the structure of modularity after removing this dominant integrating factor. This is done using the method described in Bookstein (1985). Porto *et al.* (2013) show that modularity is frequently more easily detected in matrices where the first principal component variation was removed and provide biological interpretations for these results.

### Drift

Selection is frequently invoked to explain morphological diversification, but the null hypothesis of drift being sufficient to explain current observed patterns must always be entertained. We can test the plausibility of drift for explaining multivariate diversification by using the regression method described in Ackermann and Cheverud (2002), or the correlation of principal component scores (Marroig and Cheverud 2004). Since both these tests use drift as a null hypothesis, failure to reject the null hypothesis is not evidence that selection was not involved in the observed pattern of diversification, only that the observed pattern is compatible with drift.

**DriftTest()**: Under drift, we expect that the current between group variance for many populations will be proportional to the ancestral population’s covariance structure, which is approximated by the pooled within-group covariance matrix. Conditions for the validity of these assumptions are reviewed in Prôa *et al.* (2013). Under these conditions, if *B* is the between group covariance matrix, and *W* is the within group covariance matrix, *t* is the time in number of generations and *Ne* is the effective population size, we have:

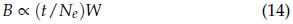

If we express all these matrices in terms of the eigenvectors of *W*, so that *W* is diagonal, we can write *B* as the variance of the scores of the means on these eigenvectors. The relationship between *B* and *W* can be expressed as a log regression, where *B*_*i*_ is the variance between groups in the projected means and 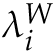 are the eigenvalues of *W*:

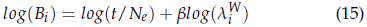

where *β* is the regression coefficient. Under drift we expect *β* to be one. If *β* is significantly different from one, we have evidence that drift is not sufficient to explain currently observed diversification.

**MultivDriftTest()**: This drift test verifies the plausibility of drift in a multivariate context when only two populations are available, one ancestral (or reference) and one derived. Let *z*_0_ represent a vector of means from *m* traits in an ancestral population. After *t* generations, the expected traits mean for *n* populations under drift would correspond to *z*_0_ with variance given by *B* = (*t*/*Ne*)*W*, where *B* represents the expected between group covariance matrix, *W* is the genetic covariance matrix from the ancestral (or reference) population, and *Ne* is the effective population size (Lande 1976, 1979; Hohenlohe and Arnold 2008). So, given the ancestral population mean and *G*-matrix, we can use this model to estimate the *B*-matrix expected under drift. We can then use this *B*-matrix as the Σ parameter in a multivariate normal distribution and sample *n* populations from this distribution. Using this sample of random populations, we can assess the amount of divergence expected by drift, estimated as the norm of the difference vectors between ancestral (or reference) and simulated population means. Then, we can compare the observed amount of divergence between the ancestral and derived populations, calculated as the norm of the difference vector between them, taking into account the standard error of traits means. An observed divergence higher than the expectations under drift indicates that genetic drift is not sufficient to explain currently observed divergence, suggesting a selective scenario.

**PCScoreCorrelation():** This test of drift relies on the correlation between principal component scores of different populations. Under drift, we expect the mean scores of different populations in the principal components of the within-group covariance matrix to be uncorrelated (Marroig and Cheverud 2004). Significant correlations between the scores of the means on any two principal components is an indication of correlated directional selection (Felsenstein 1988).

### Random matrices

**RandomMatrix():** Generating realistic random covariance matrices for null hypothesis testing is a challenging task, since random matrices must adequately sample the space of biologically plausible evolutionary parameters, like integration and flexibility. Most common covariance and correlation matrix sampling schemes fail at this, producing matrices with unrealistically low levels of integration, unless the level of integration is supplied *a priori* (as in Haber (2011)). The method described in Numpacharoen and Atsawarungruangkit (2012) provides correlation matrices with a reasonable range of evolutionary characteristics. However, the adequacy of the generated matrices in hypothesis testing has not been well established, and we recommend these random matrices be used only for informal tests requiring an arbitrary covariance or correlation matrix.

## Software availability

The most recent version of the **EvolQG** package can be installed from GitHub using the package **devtools**:

> **library** (devtools)

> **install _** github (“ lem-usp **/** evolqg “)

A less up-to-date version is also available from CRAN:

> **install**. **packages** (“ evolqg “)

## Summary

We have described a suite of functions dedicated to analyzing multivariate data sets within an evolutionary quantitative genetics framework. These functions focus on the central role that covariance and correlation matrices play in this framework; therefore, we provide functions that perform both descriptive statistics and hypothesis testing related to such matrices within an evolutionary context.

We have intentionally neglected to include techniques like phylogenetic regression or more extensive linear model functionality. We also lack Bayesian implementations that would be possible for some functions (i.e. Aguirre *et al.* (2014)). Our reasons for this are twofold: the difficulty in transposing these methods efficiently to multiple traits, especially with respect to Bayesian implementations of existing functions, and the many different robust packages for performing some of these analyses, such as phytools, phylolm, pgls, nlme, MCMCglmm and others.

Some of the material implemented here is available in other sources or through custom implementations. We have attempted to create a single consistent source for these techniques. This is by no means an exhaustive effort, and we hope to expand it given demand from the community and further developments in the field. We hope to contribute to standardization and wide adoption of these tools, and, since we opted for an open source implementation under R, this also allows the involvement of the R community in using, debugging and complementing these tools, in an effort to contribute to an open scientific environment in which, for example, truly reproducible results are the norm rather than the exception.

## Author contributions

D.M. compiled existing implementations, re-factored existing functions and contributed new code, documentation and unit tests. G.G. provided the initial set of functions and first implementations. A.H. tested and revised code and documentation. A.P.A. contributed code and documentation, G.M. developed methods and devised necessary elements for package. All authors wrote the paper.

## Competing interests

The Authors have no competing interests

## Grant information

This work was supported by funding from FAPESP. D.M. was funded by grants 2012/20180-0 and 2014/01694-9; G.G. was funded by grant 2011/52469-4; A.H. was funded by grant 2012/24937-9; A.P.A was funded by grants 2008/56886-6 and 2010/52369-0; G.M. was funded by grant 2011/14295-7.

## Acknowledgements

We would like to thank Barbara Costa, Daniela Rossoni, Edgar Zanella and Fabio Machado for contributing code and revising documentation and results.

